# Melanocore uptake by keratinocytes occurs through phagocytosis and involves Protease-activated receptor-2 activation

**DOI:** 10.1101/2021.04.13.439501

**Authors:** Hugo Moreiras, Matilde V. Neto, Liliana Bento-Lopes, Cristina Escrevente, José S. Ramalho, Miguel C. Seabra, Duarte C. Barral

## Abstract

In the skin epidermis, melanin is produced and stored within melanosomes in melanocytes and then transferred to keratinocytes. Different models have been proposed to explain the melanin transfer mechanism, which differ essentially in how melanin is transferred – either in a membrane-bound melanosome or as a melanosome core, i.e. melanocore. Here we investigated the endocytic route followed by melanocores and melanosomes during internalization by keratinocytes, by comparing the uptake of melanocores isolated from the supernatant of melanocyte cultures with melanosomes isolated from melanocytes. We show that inhibition of actin dynamics impairs the uptake of both melanocores and melanosomes. Moreover, depletion of critical proteins involved in actin-dependent uptake mechanisms, namely Rac1 and CtBP1/BARS, together with inhibition of Rac1-dependent signaling pathways or macropinocytosis suggest that melanocores are internalized by phagocytosis, whereas melanosomes are internalized by macropinocytosis. Furthermore, we confirmed that melanocore, but not melanosome uptake is dependent on the Protease-activated receptor-2 (PAR-2) and found that PAR-2 can be specifically activated by melanocores. As skin pigmentation was shown to be regulated by PAR-2 activation, our results further support the melanocore mechanism of melanin transfer and further refine this model, which can now be described as coupled melanocore exo/phagocytosis.

## Introduction

Skin pigmentation is the result of the crosstalk between melanocytes and keratinocytes. Melanocytes are highly specialized cells that synthesize the pigment melanin and contact the basal layer of the epidermis. Keratinocytes are present in all the layers of the epidermis and are the final recipients of melanin [1–4]. Melanin synthesis occurs in specialized organelles called melanosomes, within melanocytes. Melanosomes are lysosome-related organelles (LROs), since they share several features with lysosomes, like low pH (at least in early stages), the presence of lysosomal proteins and catalytic enzymes, and are secreted. Once fully mature and located at the tips of melanocyte dendrites, melanosomes are transferred to keratinocytes [5–7]. Regarding the mechanism of melanosome transfer from melanocytes to keratinocytes, there are currently four proposed models; cytophagocytosis, membrane fusion, phagocytosis of shed melanosome-laden vesicles, and coupled exo/endocytosis of the melanin core or melanocore [8–10].

The Protease-activated receptor-2 (PAR-2) is one of the few molecular players described to control melanin transfer both *in vivo* and *in vitro*. Moreover, it is known that PAR-2 stimulation leads to the enhancement of melanin transfer, as well as keratinocyte phagocytosis [11–15]. Additionally, we showed that PAR-2 regulates the uptake of melanocores, but not melanosomes [16]. Nonetheless, it is not clear what is the route followed by melanin to enter keratinocytes. The models of shed melanosome-laden vesicles and coupled exo/endocytosis of melanocores are the ones that received more attention during recent years and for which there is stronger published evidence [16,17]. Although these models are not mutually exclusive, in one the pigment is transferred as a vesicle loaded with melanosomes, and in the other as “naked” melanin. Melanosome size varies from 0.5 μm to 2 μm [18]. Therefore, phagocytosis and macropinocytosis are the internalization routes that allow the uptake of such large cargo [19,20]. Moreover, while macropinocytosis is a general process performed by most cells to uptake nutrients from the surrounding environment, phagocytosis is a process performed mainly by professional phagocytic cells and is usually specific for engulfed cargo, as it is receptor-mediated. Both these routes are highly dependent on actin cytoskeleton remodeling for phagosome and macropinosome cup formation [21]. Regarding the molecular players involved in these two processes, they are mostly shared. Indeed, both pathways are largely dependent on members of the Rho family of GTPases, namely Rho, Rac and Cdc42, which induce actin polymerization and membrane ruffling. Furthermore, Rac1 is described to be crucial for these processes during phagocytosis [22]. On the other hand, macropinosome fission from the plasma membrane is specifically dependent on Brefeldin A-ADP ribosylated substrate (CtBP1/BARS) during macropinocytosis, after cargo engulfment [23]. Additionally, the inhibition of Na+/H+ exchange by amiloride has been shown to block macropinocytosis [24].

Here we show that melanocore internalization is Rac1-dependent and CtBP1/BARS-independent, whereas melanosome uptake is CtBP1/BARS-dependent and Rac1-independent. Moreover, we observed that only melanosome internalization is significantly impaired upon treatment with an amiloride derivative, namely 5-(N-Ethyl-N-isopropyl)-Amiloride (EIPA). Furthermore, we show that melanocores, but not melanosomes are able to induce PAR-2 activation and internalization. Thus, our results further support the model of coupled exo/endocytosis and indicate that melanocore internalization occurs through phagocytosis in a PAR-2-dependent manner, with Rac1 involvement.

## Methods

### Cell culture and reagents

Mouse keratinocytes (XB2 cell line), obtained from Dorothy Bennett and Elena Sviderskaya lab, were cultured in DMEM supplemented with 10% FBS, 2 mM L-Glutamine, 100 U/ml penicillin and 100 μg/ml streptomycin. Cells were maintained in a humidified incubator at 37°C and 10% CO_2_. MNT-1 human melanoma cells were maintained in DMEM containing 10% FBS, 2 mM L-Glutamine, non-essential amino acids, sodium pyruvate, 100 U/ml penicillin and 100 μg/ml streptomycin. SLIGRL-NH2, SFLLRN and synthetic melanin were purchased from Sigma (Darmstadt, Germany), and melanin from *Sepia officinalis* from Santa Cruz (Dallas, TX, USA).

### Melanocore preparation and uptake assay

Melanocores were prepared as described previously [16]. MNT-1 cells were cultured in 150 cm^2^ flasks (Corning, NY, USA) for 5 days. Conditioned medium was collected and centrifuged at 300 × *g* for 5 minutes to pellet floating cells. The supernatant was then transferred to a clean tube and centrifuged for 1 hour and concentrated in a vivaspin Centricon (Sigma) with a pore size of 300,000 Da at 2,683 × *g*. Melanocore solution absorbance was measured using a Nanodrop 2000 (Thermo Scientific, Thermo Fisher Scientific, Waltham, MA, USA) at 340 nm and the concentration calculated according to a calibration curve developed in-house [Absorbance = 1.8546 × Concentration (g/l) – 0.0422]. For internalization studies, a concentration of 0.1 g/l of melanocores was used, unless otherwise stated.

XB2 keratinocytes were fed with 0.1 g/l of melanocores or melanosomes and incubated at 37°C and 10% CO_2_ for 24h. Cells were then washed 3× with PBS and fixed with 4% paraformaldehyde (PFA) (Electron Microscopy Sciences, PA, USA) in PBS for 20 minutes at room temperature. The number of melanocores per cell was counted and the mean calculated. Nuclei were counted to ensure similar cell confluence in all samples and the amount of melanin internalized per cell calculated. In the assays using inhibitors, cells were incubated with EIPA (Tocris, MN, USA; 50 μM) or EHT 1864 (Santa Cruz Biotechnology, Dallas, TX, USA; 20 μM) for 20 minutes or overnight, respectively, before being fed with melanocores (0.1 g/l), melanosomes or Rhodamine B isothiocyanate–Dextran 70kDa (Sigma, Darmstadt, Germany; 0.1μg/μl) for 2h and processed for immunofluorescence as described below. In the experiments with dextran, melanin-fed cells were stained with anti-PMEL antibody (HMB45) (Agilent DAKO, CA, USA), to allow the detection of internalized melanin by confocal microscopy and quantification using the Spot Detector plugin from Icy Software (https://icy.bioimageanalysis.org).

### Melanosome preparation

Melanosomes were prepared as described previously (Chabrillat et al. 2005). Briefly, MNT-1 cells were scraped in H buffer (50 mM imidazole, pH 7.4, 250 mM sucrose, 1 mM EDTA, 0.5 mM EGTA, 5 mM MgSO_4_, 0.15 mg/ml casein and 1 mM dithiothreitol and homogenized by passing several times through a cell cracker 26G needle. The post-nuclear supernatant (PNS) was obtained by centrifugation at 600 × *g* for 5 minutes. The PNS was then centrifuged at 2,500 × *g* for 5 minutes. The recovered pellet was added onto a 50% Percoll cushion, centrifuged at 5,000 × *g* for 20 minutes, and the pellet of melanosomes collected.

### Immunofluorescence microscopy

Cells were grown on coverslips for immunofluorescence and fixed in 4% PFA in PBS for 20 minutes at room temperature. Cells were blocked and permeabilized with 1% bovine serum albumin (Sigma) and 0.05% saponin (Sigma) in PBS for 30 minutes. Fixed cells were then incubated with primary antibodies for 1 hour and then for a further 1 hour with appropriate secondary antibodies conjugated with a fluorophore (Molecular Probes). The antibodies used and respective dilutions are displayed in Supplementary Table 1. Coverslips were finally mounted in MOWIOL mounting medium (Calbiochem). All antibody incubations and washes were done with 1 × PBS, 0.5% BSA and 0.05% saponin. To visualize the nuclei, cells were incubated with DAPI (Invitrogen) for 5 minutes. Images were acquired on a Zeiss Observer Z2 widefield microscope, equipped with a Zeiss 506 Mono camera using the 63× 1.4 NA Oil objective or a Zeiss LSM 710 confocal microscope with a Plan-Apochromat 63× 1.4 NA oil-immersion objective. Images were processed using ImageJ and Adobe Illustrator 6.0 (Adobe, San Jose, CA, USA) software.

### Flow cytometry

To prepare XB2 keratinocytes for flow cytometry, cells were washed with PBS and FACS buffer [1% FBS (v/v) and 2 mM EDTA in PBS], centrifuged at 300 × *g* for 5 minutes and resuspended in 200 μl FACS buffer. Cells were incubated with anti-FLAG-Cy3 (Sigma) antibody, diluted 1:50 in FACS buffer for 1 h, at 4°C to ensure that internalization was blocked. Cells were then washed with FACS buffer, centrifuged at 300 × *g* for 5 minutes and resuspended with 200 μl FACS buffer. Data acquisition was performed in a FACS CANTO II flow cytometer (BDBiosiences) At least 30,000 cells were acquired per condition using BD FACSDivaTM software (Version 6.1.3, BD Biosciences). Data analysis was performed in FlowJo (Version 10, BD Biosciences).

### Actin polymerization inhibition assay

XB2 keratinocytes (5 × 10^4^) were treated with 0.02 μg/ml Lantrunculin A (Sigma) or 0.05 μg/ml Cytochalasin D (Sigma), both diluted in DMSO or non-treated for 1 hour and then incubated at the same time with melanocores for 24 hours. Cells where then washed and left to recover for 1 hour in complete growth medium before fixation with 4% (v/v) PFA in PBS. Nuclei were visualized by DAPI staining, and filamentous (F)-actin was stained with Phalloidin 568.

### Plasmid transfection

XB2 mouse keratinocytes (4 × 10^4^) cultured in 24-well plates 1 μg of pEGFP-FLAG-PAR-2-HA diluted in 100 μl of Opti-MEM were mixed with 1.5 μl of TurboFect (Thermo Scientific) in 100 μl of Opti-MEM according to the manufacturer’s instructions. Cells were incubated for 24 hours at 37°C and then the medium was changed to 500 μl complete XB2 medium.

### siRNA silencing

XB2 keratinocytes (1 × 10^5^ per well) were seeded in 24-well plates (Corning – Corming, NY, USA). Twenty-four hours later, 50 nM of gene-specific siGenome SMART pool (Thermo Scientific) were diluted in 32 μl Opti-MEM (Gibco), while 1.2 μl of Dharmafect 1 (Dharmacon, GELifeScience) were added to 6 μl of Opti-MEM. These mixtures were combined and incubated at room temperature for 20 minutes. Finally, 160 μl of Opti-MEM were added to the mixture, the medium was removed from the cells, and the siRNA mixture added to each well. Cells were incubated for 24 hours at 37°C and then the medium was changed to complete medium. Non-targeting siRNA pool (Thermo Scientific) was used as control.

### Real-time quantitative PCR

Total RNA was isolated from cells using RNeasy Mini kit (Qiagen) and reverse-transcribed to cDNA using SuperScript^®^ II (Invitrogen), according to the manufacturer’s instructions. qRT-PCR reactions were performed using a Roche LightCycler equipment (Roche) and Roche SybrGreen Master Mix reagent (SybrGreen, Roche). Five μl of SybrGreen, 4 μl of cDNA together with 10 μM of appropriate primers were used per well, in triplicate, for a total reaction volume of 10 μl. For each protein, gene expression was calculated relative to control wells and normalized for *GAPDH* (used as housekeeping gene), using LightCycler96 software (Roche) to analyze the results. The primers used are shown in Supplementary Table 2.

### Statistical analysis

Statistical analysis was performed using Prism software (GraphPad Software Inc.). Unpaired Student’s t-test was used to analyze the *p* value of the differences. Colocalization in microscopy images was measured using Pearson’s correlation coefficient with ImageJ software and the plugin JACoP (Just Another Colocalization Plugin).

## Results

### Melanocore uptake is dependent on actin polymerization

To determine the mechanism by which melanin is internalized, we started by probing for the dependency on actin by using the actin polymerization inhibitory drugs cytochalasin D and latrunculin A. To ingest particles by phagocytosis and extracellular fluid by macropinocytosis, cells form plasma membrane protrusions that close at their distal end, giving rise to membrane-bound organelles termed phagosomes or macropinosomes, respectively [22,23]. Due to the size of the protrusions formed during these internalization processes, actin cytoskeleton remodeling is essential to reshape and extend the plasma membrane. Since melanosome size varies from 0.5 μm to 2 μm, it is likely that melanin is internalized through phagocytosis or macropinocytosis [18]. Therefore, actin polymerization inhibition is expected to impair melanin internalization. To test this hypothesis, we treated XB2 mouse keratinocytes with latrunculin A or cytochalasin D and then incubated the cells with melanocores isolated from the culture medium of MNT-1 mouse melanoma cells. We found that both cytochalasin D and latrunculin A lead to a ~30% reduction in the number of melanocores internalized by keratinocytes, when compared with non-treated cells or cells treated with DMSO (Figure 1). Additionally, we found that the internalization of melanosomes isolated from MNT-1 melanocytes is also dependent on actin polymerization (Figure 2), since the treatment with cytochalasin D or latrunculin A results in a 30% reduction in internalized melanin, relative to control cells. Therefore, these results show that melanin uptake is actin-dependent, regardless of the form of melanin presented to the cells.

**Figure 1.**
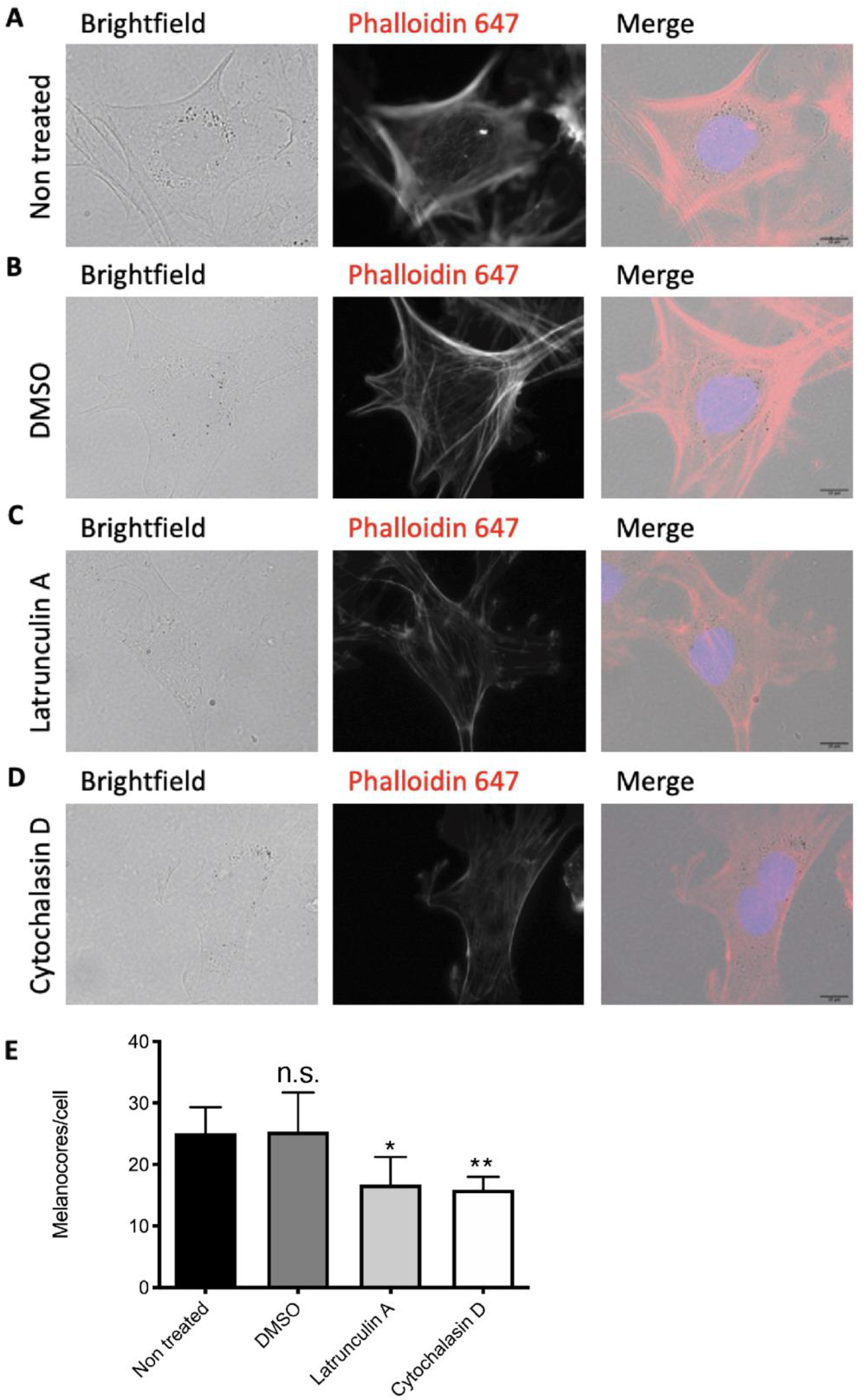
Melanocore uptake is actin-dependent. XB2 mouse keratinocytes were (A) left untreated or treated with (B) DMSO (vehicle), (C) Latrunculin A, or (D) Cytochalasin D for 1h and then incubated with melanocores for 24h. Cells where washed and allowed to recover for 1h before fixation. Nuclei were visualized by DAPI staining (blue) and F-actin stained with Phalloidin 647 (red). Scale bars = 10 μm. (E) Quantification of melanocore number per cell. *p* values (unpaired Student t-test) were considered statistically significant when <0.05 (*), <0.01 (**) or non-significant (n.s.) when >0.05. Plots show mean ± SD of three independent experiments.

**Figure 2.**
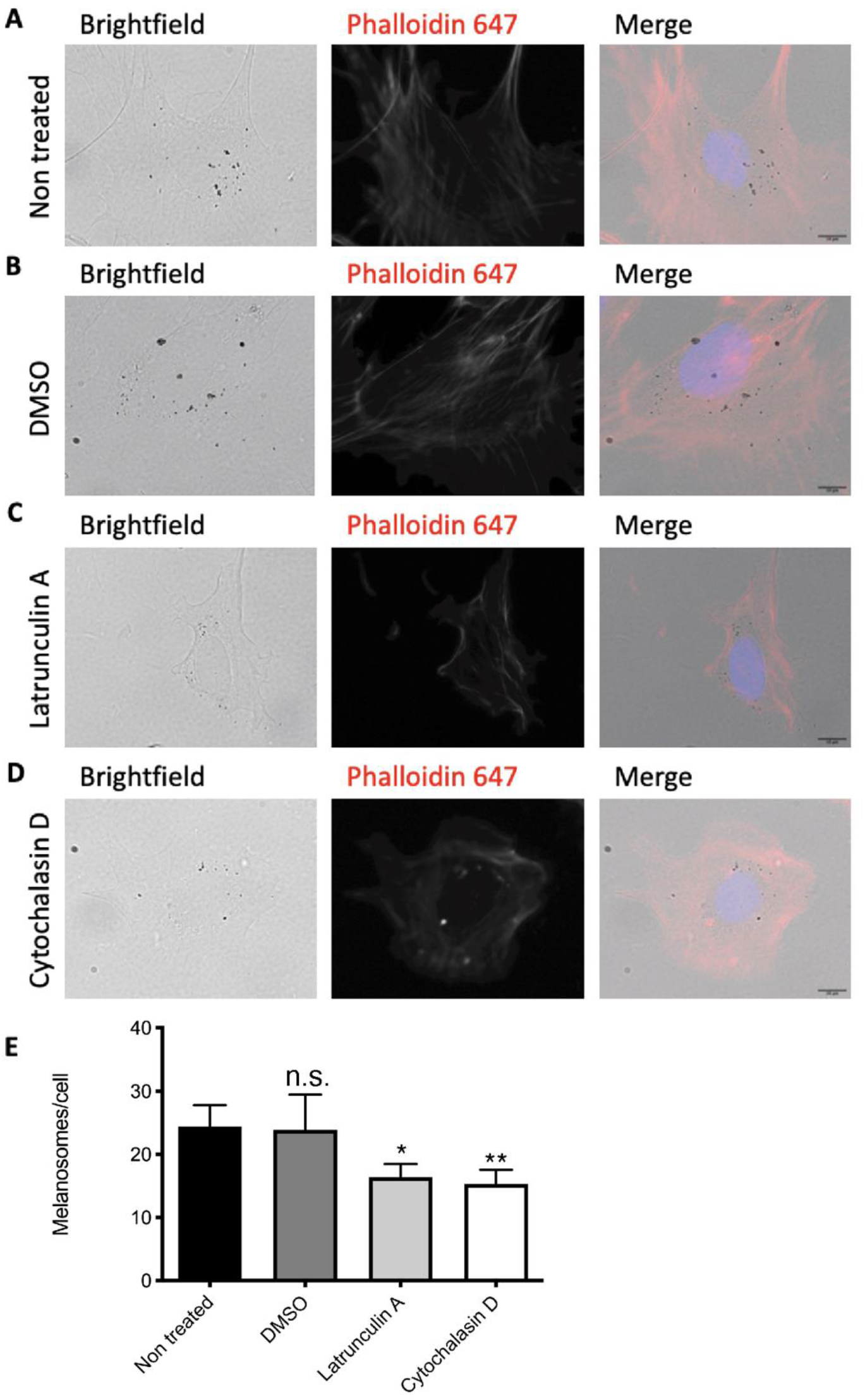
Melanosome uptake is actin-dependent. XB2 mouse keratinocytes were (A) left untreated or treated with (B) DMSO (vehicle), (C) Latrunculin A, or (D) Cytochalasin D for 1h and then incubated with melanosomes for 24h. Cells where washed and allowed to recover for 1h before fixation. Nuclei were visualized by DAPI staining (blue) and F-actin stained with Phalloidin 647 (red). Scale bars = 10 μm. (E) Quantification of melanosomes number per cell. *p* values (unpaired Student t-test) were considered statistically significant when <0.05 (*), <0.01 (**) or non-significant (n.s.) when >0.05. Plots show mean ± SD of three independent experiments.

### Melanocore internalization is impaired by Rac1 interference but not macropinocytosis blockade

Phagocytosis and macropinocytosis play key roles during development and although they are essential mechanisms of internalization, there are important differences between them. Phagocytosis allows the internalization of large extracellular particles, while macropinocytosis is used by cells to uptake extracellular fluid and soluble cargo from the surrounding environment [19,20]. Regarding the molecular players involved in these two processes, they overlap significantly, since both are largely dependent on the recruitment of cortical actin to the plasma membrane for cargo engulfment. The major difference is that macropinocytosis is a general process performed by the majority of the cells to uptake nutrients from the extracellular milieu, while phagocytosis is mainly performed by specialized cells and is usually specific for the particles engulfed as it is receptor-mediated. Nevertheless, keratinocytes were shown to possess phagocytic ability [14,25,26]. Rac1 and CtBP1/BARS are essential for actin polymerization and membrane ruffling in phagocytosis, as well as the fission step of macropinosome formation during macropinocytosis, respectively [22,23,27]. To characterize the endocytic route followed by melanin, we silenced Rac1 or CtBP1/BARS, in XB2 keratinocytes (Supplementary Figure 1), before incubating them for 24h with melanocores or melanosomes. We found that Rac1 silencing impairs melanocore uptake by 82% when compared with control cells, whereas CtBP1/BARS depletion does not affect melanocore internalization (Figure 3). As expected, melanocore, but not melanosome internalization is dependent on PAR-2, confirming our previous studies (Figure 3A and B and Figure 4 A and B) [16]. In the case of melanosomes, we observed that CtBP1/BARS, but not Rac1 depletion reduces internalization by approximately 35% (Figure 4A and B). Moreover, we used the Rac1 inhibitor EHT 1864 and observed a decrease of about 55% in melanocore uptake upon treatment with this drug (Figure 3C and D). However, no differences were observed in melanosome internalization upon treatment with EHT 1864 (Figure 4C and D). Furthermore, we treated XB2 keratinocytes with the macropinocytosis inhibitor EIPA and assessed internalization of melanosomes or melanocores. We observed that while melanocore internalization is not affected by this drug, melanosome internalization is decreased by approximately 50% (Figure 3C and D and Figure 4C and D). These results suggest that melanosomes are internalized by macropinocytosis, whereas melanocore internalization has the characteristics of a phagocytic process.

**Figure 3.**
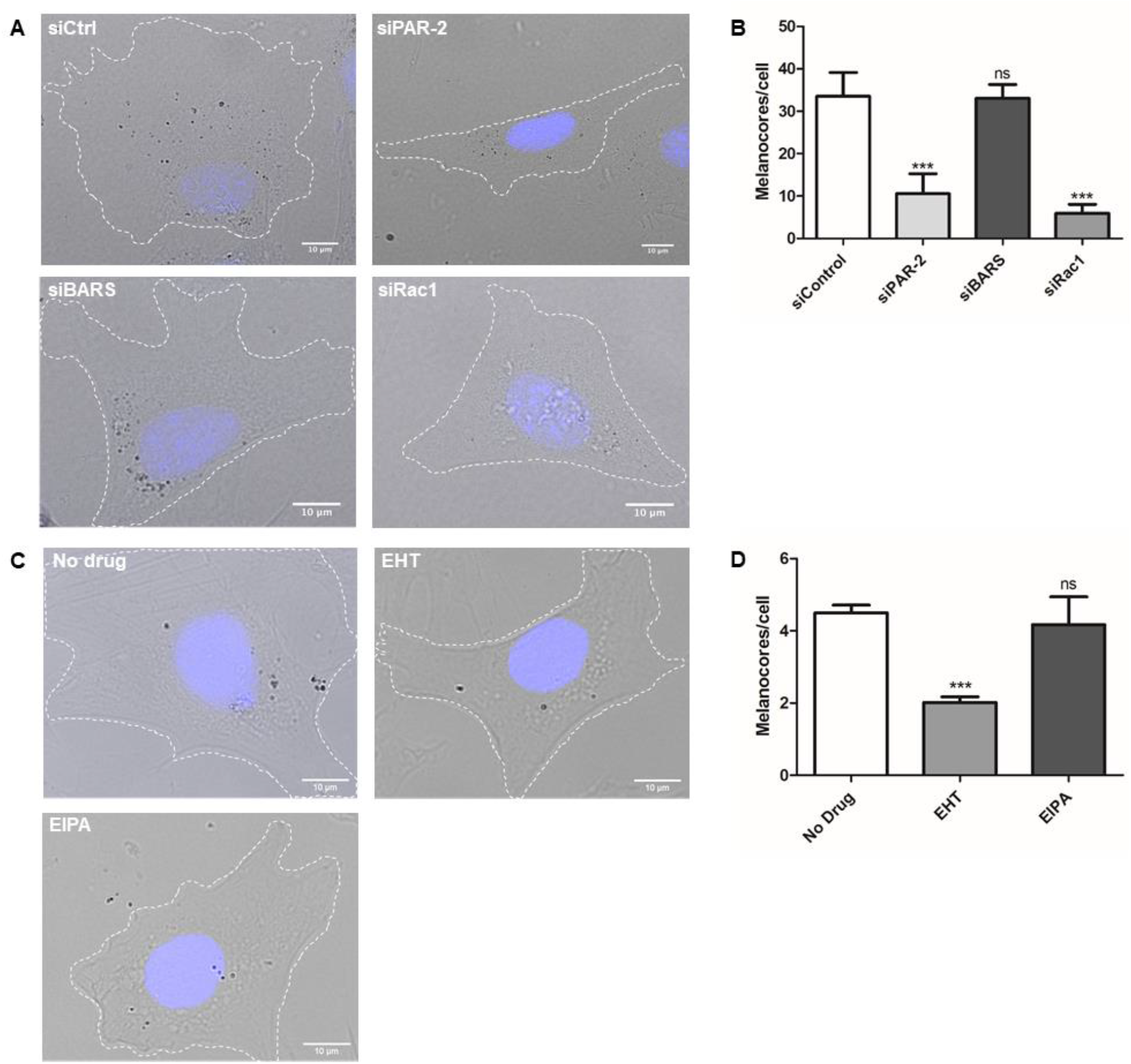
Melanocore uptake is PAR-2- and Rac1-dependent. XB2 keratinocytes were transfected with non-targeting siRNA (siCtrl), siRNA targeting PAR-2 (siPAR-2), Rac1 (siRac1) or BARS (siBARS) (A and B) and treated or not with EHT 1864 (EHT) or EIPA (C and D). Cells were then incubated with 0.1 g/l of melanocores. The transfected cells and the cells treated with the drugs were fixed 24 hours or 2 hours later, respectively and examined by brightfield microscopy. Nuclei were visualized by DAPI staining (blue). Scale bars = 10 μm. (B) Quantification of internalized melanocores/cell after a 24h pulse. (D) Quantification of internalized melanocores/cell after a 2h pulse. *p* values (unpaired Student t-test) were considered statistically significant when <0.05 (*), <0.01 (**), <0.001 (***) or non-significant (n.s.) when >0.05. Plots show mean ± SD of three independent experiments.

**Figure 4.**
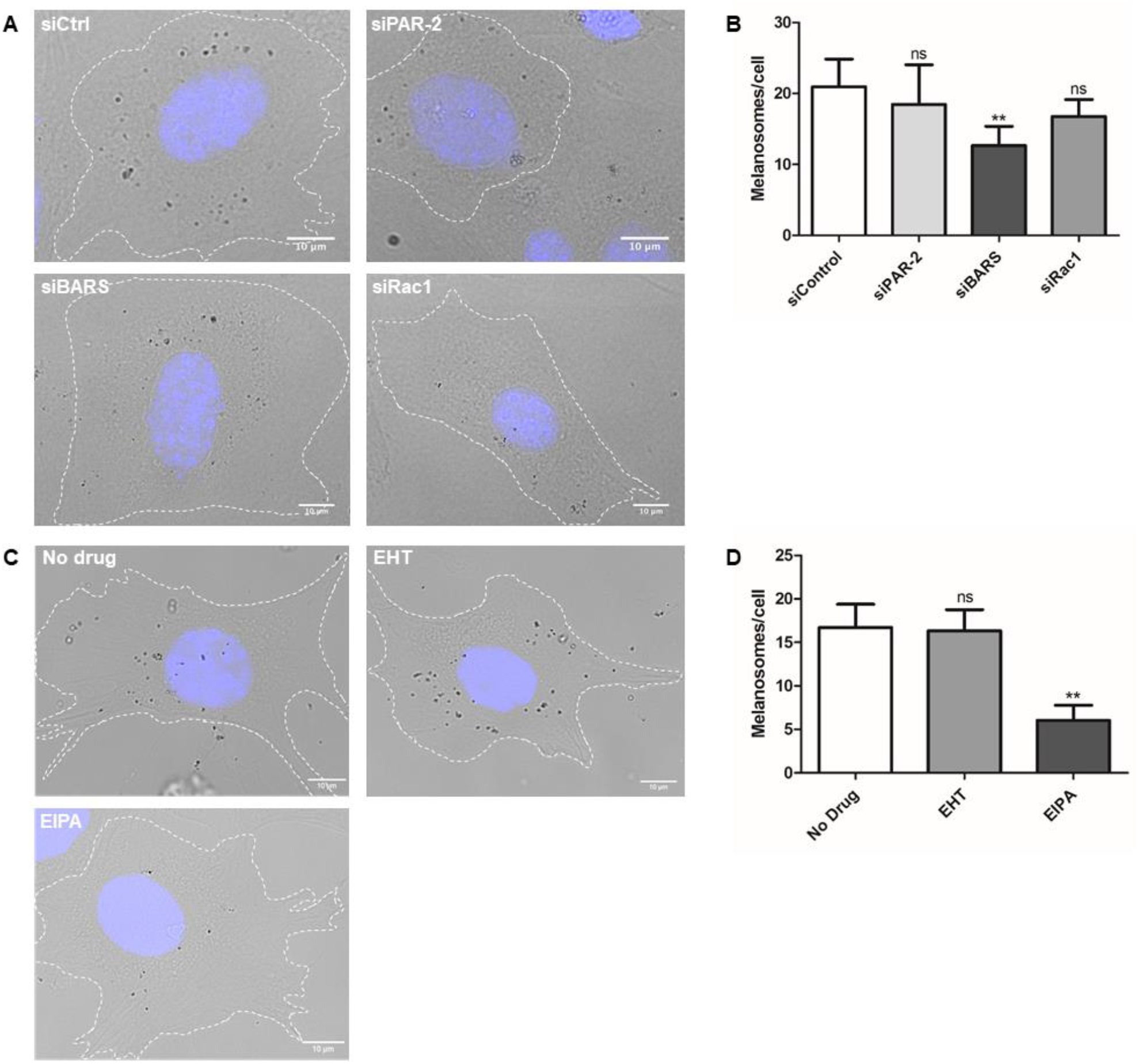
Melanosome uptake is CtBP1/BARS-dependent. XB2 keratinocytes were transfected with non-targeting siRNA (siCtrl), siRNA targeting PAR-2 (siPAR-2), Rac1 (siRac1) or BARS (siBARS) (A and B) and treated or not with EHT 1864 (EHT) or EIPA (C and D). Cells were then incubated with 0.1 g/l of melanosomes. The transfected cells and the cells treated with the drugs were fixed 24 hours or 2 hours later, respectively and examined by brightfield microscopy. Nuclei were visualized by DAPI staining (blue). Scale bars = 10 μm. (B) Quantification of internalized melanosomes/cell after a 24h pulse. (D) Quantification of internalized melanosomes/cell after a 2h pulse. *p* values (unpaired Student t-test) were considered statistically significant when <0.05 (*), <0.01 (**), <0.001 (***) or non-significant (n.s.) when >0.05. Plots show mean ± SD of three independent experiments.

To confirm these results, we used as a positive control 70 kDa dextran, which was shown to be internalized predominantly via macropinocytosis in an amiloride-sensitive manner [28]. In this case, we quantified melanin internalization by immunofluorescence, staining melanin-containing compartments in keratinocytes with an antibody against PMEL, an essential structural protein in melanosome biogenesis. Importantly, both melanocores and melanosomes are stained within keratinocytes using this antibody (Supplementary Figure 2A), allowing the accurate quantification of melanin-containing compartments. Strikingly, EIPA treatment reduces melanosome internalization by 55%, similar to the decrease observed for dextran (~56%) (Supplementary Figure 2B). As expected, CtBP1/BARS depletion significantly impairs dextran internalization (~32%), confirming that it occurs via macropinocytosis (Supplementary Figure 2C). In contrast, we found that melanocore uptake is almost unaffected by EIPA treatment (Supplementary Figure 2C), further suggesting that melanocores are not internalized by macropinocytosis. Altogether, these results suggest that melanocore internalization occurs through phagocytosis, whereas melanosomes are internalized through macropinocytosis.

### Melanocores but not melanosomes induce PAR-2 internalization

PAR-2 is a transmembrane receptor with an extracellular amino‐terminus motif that acts as an activating ligand upon receptor cleavage. Interestingly, PAR-2 is expressed in keratinocytes but not in melanocytes and it is thought that the increase in melanin transfer and internalization that occurs upon PAR-2 activation is due to an increase in actin dynamics [11,14,15]. To elucidate the role of PAR-2 in melanin uptake, we assessed the receptor internalization upon incubation with melanocores, melanosomes or activating/control peptides (SLIGRL/SFLLRN, respectively). For this, XB2 keratinocytes were transfected with a plasmid encoding PAR-2, FLAG-tagged at the N-terminus and HA-tagged at the C-terminus. After stimulation and internalization of the receptor, the N-terminus is cleaved, and the FLAG-tag is lost. Indeed, after 24h of incubation with SLIGRL peptide, which is known to activate PAR-2, the levels of FLAG at the plasma membrane decrease significantly, by approximately 20%, when compared to non-stimulated cells or cells incubated with the control peptide SFLLRN (Figure 5B and C). These results were quantified by flow cytometry by measuring the mean fluorescence intensity (MFI) of FLAG at the cell surface, and by immunofluorescence through the analysis of the Pearson’s correlation coefficient between FLAG and HA tags at the plasma membrane (Figure 5F and G). Interestingly, we observed that melanocores but not melanosomes can activate PAR-2 and induce its internalization by approximately 20% (Figure 5D-G). Therefore, these observations suggest that melanocores but not melanosomes activate PAR-2, similar to the activating peptide. Moreover, our results confirm that melanocore but not melanosome internalization is mediated by PAR-2.

**Figure 5.**
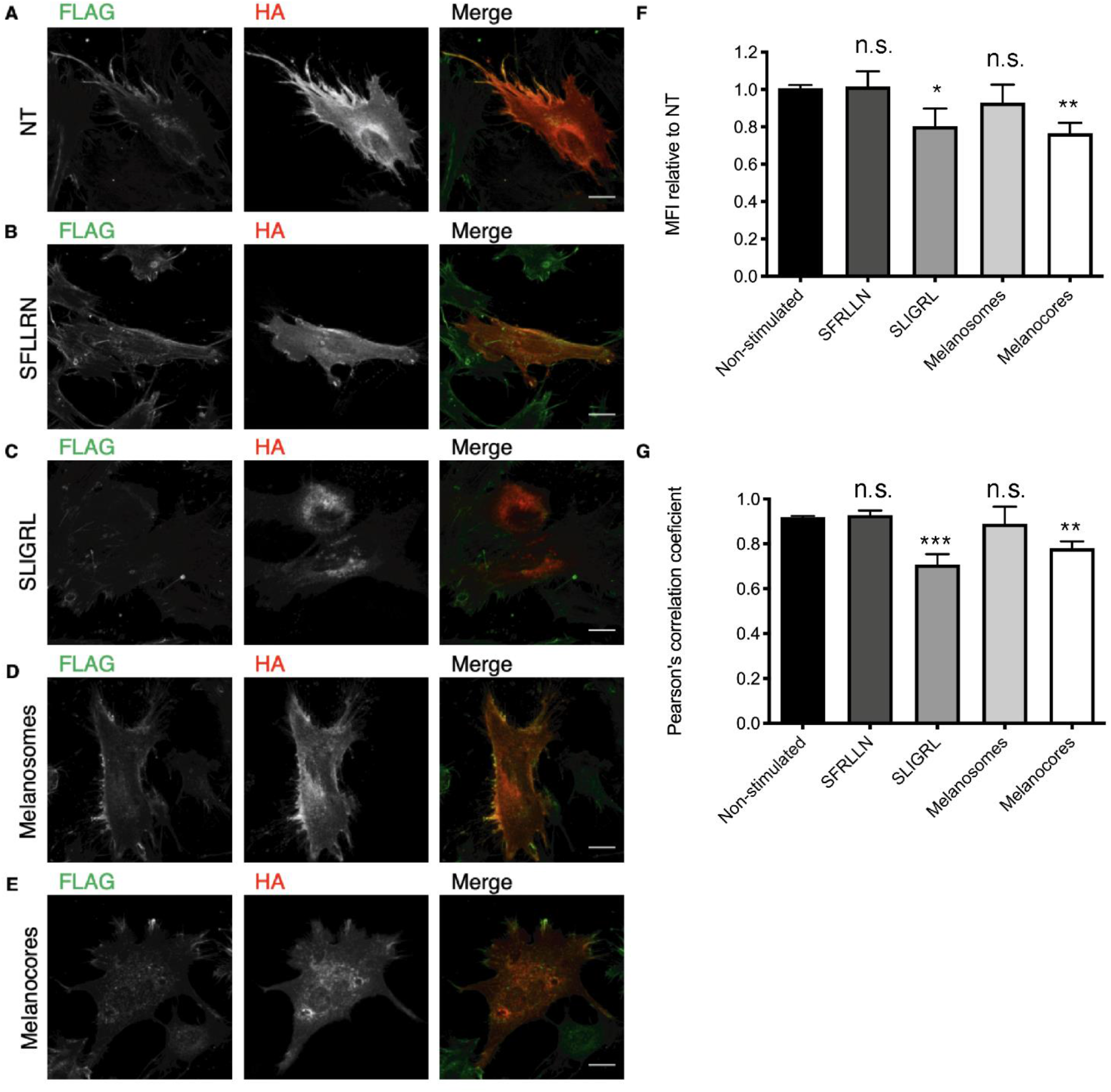
Melanocores but not melanosomes induce PAR-2 internalization. XB2 keratinocytes were transfected with FLAG-PAR-2-HA and after 24h were stimulated or not (NT) (A) with SFRLLN (B), SLIGRL (C), Melanosomes (D) or Melanocores (E). After 24h, cells were either fixed and stained for FLAG (green) and HA (red) tags or processed for flow cytometry and stained with FLAG antibody Cy3-conjugated to assess the levels of FLAG at the cell surface. (A-E) Representative images of XB2 keratinocytes after 24h stimulation. Scale bars = 10 μm. (F) Quantification of FLAG cell surface levels after 24h stimulation. (G) Graphical representation of Pearson’s coefficient between FLAG (green) and HA (red) pixels at cell periphery. *p* values (unpaired Student t-test) were considered statistically significant when <0.05 (*), <0.01 (**), <0.001 (***) or non-significant (n.s.) when >0.05. Plots show mean ± SD of three independent experiments.

## Discussion

We reported that melanocores but not melanosomes require PAR-2 to be internalized, suggesting that PAR-2 activation has specificity for non-membrane-bound melanin [16]. Here we further demonstrate that melanocores and melanosomes are internalized by keratinocytes through distinct routes and that only melanocores are internalized by a PAR-2-dependent phagocytic process.

Whilst we observed that both melanocores and melanosomes are dependent on actin to be internalized, we present evidence suggesting that melanocore internalization is mediated by phagocytosis and melanosome uptake occurs by macropinocytosis. To distinguish between these two processes, we depleted Rac1 or CtBP1/BARS, which are important regulators of the aforementioned internalization routes [22,23], and quantified melanocore and melanosome internalization. We found that melanocore internalization is dependent on Rac1, in contrast with melanosomes, which are internalized in a CtBP1/BARS-dependent and Rac1-independent manner. Furthermore, Rac inhibition using EHT 1864 significantly decreases melanocore uptake, without affecting the number of melanosomes internalized by keratinocytes. Similar results were obtained upon Rac1 depletion. Rac1 activation and membrane recruitment was described to have an important role in actin polymerization and membrane remodeling during the formation of the phagocytic cup [29], therefore regulating the early steps of particle internalization. Although Rac1 activation was also shown to be sufficient to induce macropinocytic cup formation, its inactivation is crucial for macropinosome closure and recruitment of molecules that regulate macropinosome maturation [30]. On the other hand, whilst melanocore internalization is not affected by macropinocytosis inhibition with EIPA, melanosome uptake is highly sensitive to the treatment with this drug. Similar to melanosomes, the internalization of 70 kDa dextran, which is described to be internalized by macropinocytosis [28] is impaired by both CtBP1/BARS depletion or EIPA treatment. Taken together, our results show that melanocore internalization is highly dependent on the presence and activation of Rac1 and unaffected by EIPA treatment, strongly suggesting phagocytosis as the route taken to enter keratinocytes. Conversely, neither the absence nor the inactivation of Rac1 impair melanosome uptake. Instead, CtBP1/BARS depletion or EIPA treatment were found to impair melanosome internalization, reinforcing the involvement of macropinocytosis in melanosome uptake.

We also assessed the ability of melanocores and melanosomes to induce PAR-2 activation and internalization. Only melanocores can activate and induce endocytosis of the receptor, suggesting a mechanism in which melanocores are internalized by phagocytosis in a Rac1-dependent manner upon activation of PAR-2, as opposed to intact melanosomes, which are internalized through macropinocytosis independently of PAR-2. Thus, PAR-2 could be essential to prepare for the formation of the phagocytic cup that engulfs melanocores during their internalization by keratinocytes.

How melanin is presented to keratinocytes, *i.e.*, the presence or absence of membrane surrounding the melanin core is therefore critically important to define the route of internalization. We speculate that melanocore transfer through coupled exo/phagocytosis occurs preferentially at basal levels of pigmentation and is the main mode of transfer, while the transfer of shed melanosome-laden vesicles could be used in stress conditions, for instance during tanning to ensure a high throughput. Indeed, a recently published study found that the uptake of melanosome-rich globules under Toll-like receptor-3 (TLR-3)-stimulated conditions is dependent on RhoA activation but independent of PAR-2 [31]. PAR-2-stimulation catalyzes the conversion of ATP to cAMP, which activates Rac1 and subsequently inactivates RhoA [15,32]. This PAR-2-mediated activation of RhoA/Rac1 may thus be a critical factor in melanin uptake. Nevertheless, the fact that TLR-3 activation leads to an increase in PAR-2 expression might explain the increase in phagocytic activity of keratinocytes, as previously reported [15,25]. Therefore, future studies should analyze the role of other Rho GTPases, including RhoA and Cdc42 in melanosome and melanocore uptake. Additionally, it is important to determine the activation status of these proteins after keratinocyte incubation with melanosomes or melanocores. In conclusion, our current results and previously published studies suggest that the main mechanism of pigment transfer in the skin epidermis involves coupled exo/phagocytosis of melanocores regulated by PAR-2.

## Acknowledgements

We would like to thank the scientific and technical assistance from the CEDOC Cell Culture, Flow Cytometry and Microscopy facilities. We also thank Dorothy Bennett and Elena Sviderskaya (St. George’s University of London, UK), Susana Lopes (CEDOC, NOVA Medical School) and Paulo Matos (National Health Institute Doutor Ricardo Jorge, Portugal) for the kind gift of reagents and Paulo Matos also for expert advice. This project was supported by Fundação para a Ciência e a Tecnologia (FCT), Portugal through grant PTDC/BIA-CEL/29765/2017, PhD fellowships to HM, MVN and LBL (PD/BD/114118/2015, PD/BD/137442/2018 and SFRH/BD/131938/2017, respectively) and the FCT Investigator Program to DCB (IF/00501/2014/CP1252/CT0001).

## Supplementary Information

**Supplementary Figure 1.**
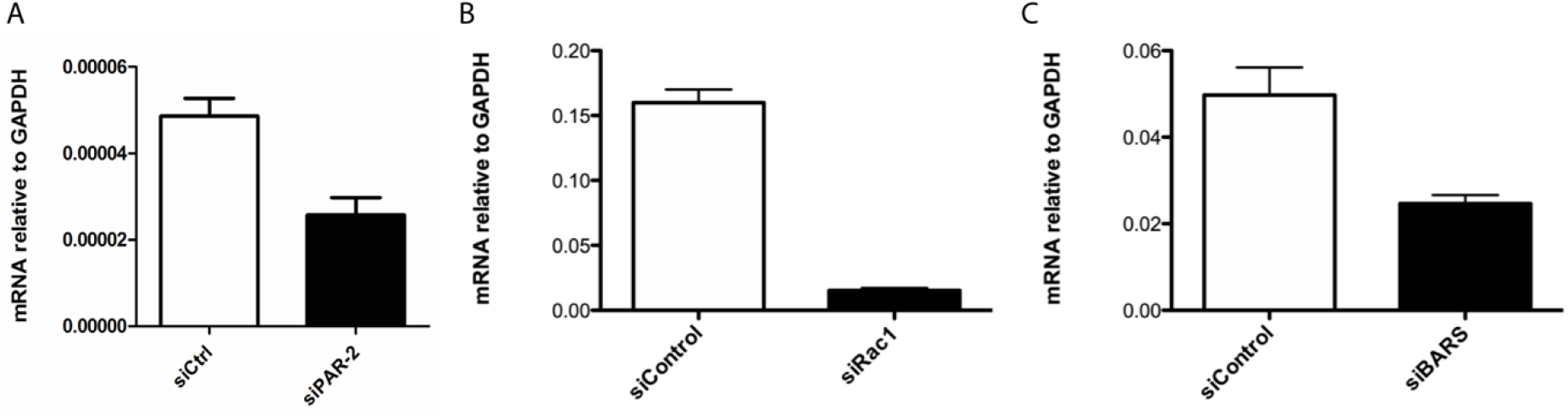
Efficiency of gene expression silencing. XB2 mouse keratinocytes were transfected with siRNAs targeting: PAR-2 (A), Rac1 (B) or BARS (C) and the levels of silencing measured by qRT-PCR after 48 hours.

**Supplementary Figure 2.**
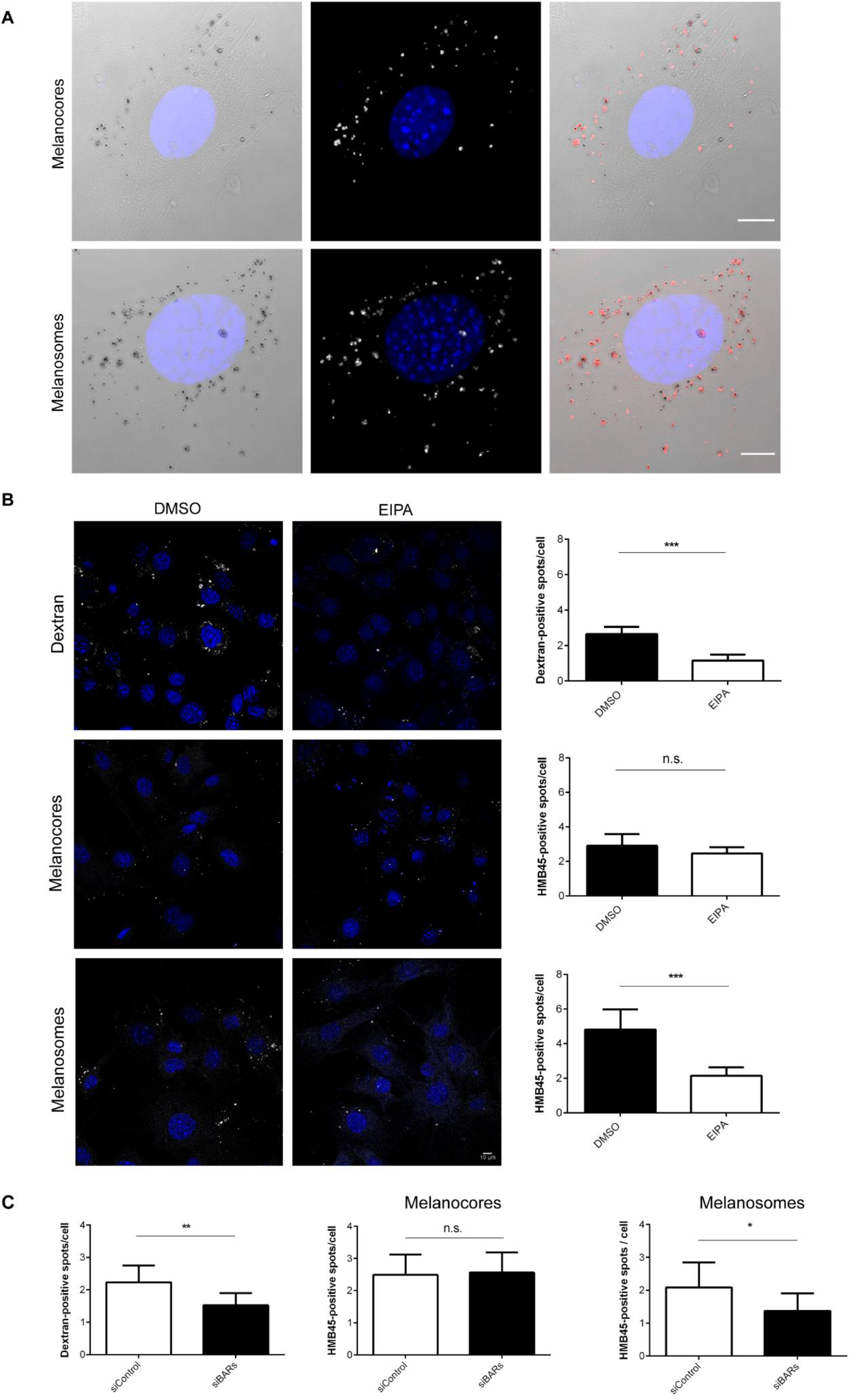
Macropinocytosis blockade impairs dextran and melanosome uptake. (A) Confocal microscopy images showing melanocores and melanosomes labeled with HMB45 antibody, which recognizes PMEL. BF, bright field. (B and C) XB2 keratinocytes were treated with EIPA or DMSO (vehicle) for 20 minutes (B) or transfected with non-targeting siRNA (siCtrl) or siRNA targeting CtBP1/BARS (siBARS) for 2 days (C) before being incubated with melanocores (0.1 g/l), melanosomes (0.1 g/l) or 70 kDa dextran (0.1 μg/μl) for 2 hours and then fixed. Quantification of internalized dextran, melanocores and melanosomes was done using the Spot Detector plugin (Icy software) and represented as the number of positive spots/cell. Scale bars = 10 μm. *p* values (unpaired Students t-test) were considered statistically significant when <0.05 (*), <0.01 (**), <0.001 (***) or non-significant (n.s.) when >0.05. Plots show mean ± SD of three independent experiments.

**Supplementary Table 1.** Antibodies and respective dilutions used in this study.

| **Antibody** | **Species** | **Brand** | **Dilution** |
| --- | --- | --- | --- |
| anti-FLAG | Mouse | Sigma F3165 | IF 1:200 |
| anti-FLAG-Cy3 | Mouse | Sigma A9594 | FC 1:50 |
| anti-HA | Rabbit | Abcam Ab9110-100 | IF 1:150 |
| Anti-HMB45 | Mouse | Dako M0634 | IF 1:200 |
| Alexa Fluor 488-anti-mouse | Donkey | Invitrogen<br>#A-11001 | IF 1:400 |
| Alexa Fluor 568-anti-rabbit | Goat | Invitrogen<br># A-21206 | IF 1:400 |

IF – Immunofluorescence, FC – Flow cytometry

**Supplementary Table 2.**
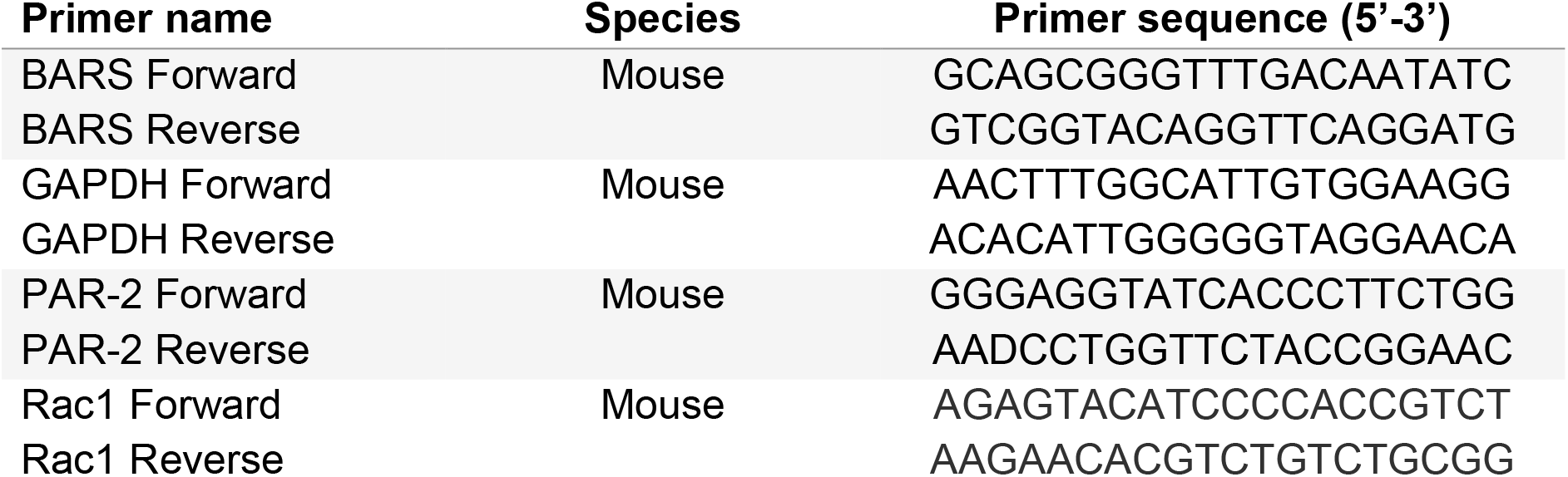
Primers used in this study.

## References

1. Lin, J.Y.; Fisher, D.E. Melanocyte biology and skin pigmentation. Nature 2007, 445, 843–850, doi:10.1038/nature05660.

2. Lai-Cheong, J.E.; McGrath, J.A. Structure and function of skin, hair and nails. Med. (United Kingdom) 2013, 41, 317–320, doi:10.1016/j.mpmed.2013.04.017.

3. Nicol, N.H. Anatomy and physiology of the skin. Dermatol. Nurs. 2005, 17, 62, doi:10.4324/9780203450505_chapter_1.

4. Marks, M.S.; Seabra, M.C. The melanosome: membrane dynamics in black and white. Nat. Rev. Mol. Cell Biol. 2001, 2, 738–48, doi:10.1038/35096009.

5. Wasmeier, C.; Hume, A.N.; Bolasco, G.; Seabra, M.C. Melanosomes at a glance. J. Cell Sci. 2008, 121, 3995–3999, doi:10.1242/jcs.040667.

6. Raposo, G.; Marks, M.S. Melanosomes — dark organelles enlighten endosomal membrane transport. Nat. Rev. Mol. Cell Biol. 2007, 8, 786–797, doi:10.1038/nrm2258.

7. Delevoye, C.; Marks, M.S.; Raposo, G. Lysosome-related organelles as functional adaptations of the endolysosomal system. Curr. Opin. Cell Biol. 2019, 59, 147–158, doi:10.1016/j.ceb.2019.05.003.

8. Wu, X.; Hammer, J.A. Melanosome transfer: It is best to give and receive. Curr. Opin. Cell Biol. 2014, 29, 1–7, doi:10.1016/j.ceb.2014.02.003.

9. Van Den Bossche, K.; Naeyaert, J.M.; Lambert, J. The quest for the mechanism of melanin transfer. Traffic 2006, 7, 769–778, doi:10.1111/j.1600-0854.2006.00425.x.

10. Yoshida, Y.; Hachiya, A.; Sriwiriyanont, P.; Ohuchi, A.; Kitahara, T.; Takema, Y.; Visscher, M.O.; Boissy, R.E. Functional analysis of keratinocytes in skin color using a human skin substitute model composed of cells derived from different skin pigmentation types. FASEB J. 2007, 21, 2829–39, doi:10.1096/fj.06-6845com.

11. Seiberg, M.; Paine, C.; Sharlow, E.; Andrade-Gordon, P.; Costanzo, M.; Eisinger, M.; Shapiro, S.S. The protease-activated receptor 2 regulates pigmentation via keratinocyte-melanocyte interactions. Exp. Cell Res. 2000, 254, 25–32, doi:10.1006/excr.1999.4692.

12. Seiberg, M.; Paine, C.; Sharlow, E.; Eisinger, M.; Shapiro, S.S.; Andrade-Gordon, P.; Costanzo, M. Inhibition of Melanosome Transfer Results in Skin Lightening1. J. Invest. Dermatol. 2000, 115, 162–167, doi:10.1046/j.1523-1747.2000.00035.x.

13. Lin, C.B.; Chen, N.; Scarpa, R.; Guan, F.; Babiarz-Magee, L.; Liebel, F.; Li, W.-H.; Kizoulis, M.; Shapiro, S.; Seiberg, M. LIGR, a protease-activated receptor-2-derived peptide, enhances skin pigmentation without inducing inflammatory processes. Pigment Cell Melanoma Res. 2008, 21, 172–183, doi:10.1111/j.1755-148X.2008.00441.x.

14. Sharlow, E.R.; Paine, C.S.; Babiarz, L.; Eisinger, M.; Shapiro, S.; Seiberg, M. The protease-activated receptor-2 upregulates keratinocyte phagocytosis. J. Cell Sci. 2000, 113, 3093–3101.

15. Scott, G.; Leopardi, S.; Parker, L.; Babiarz, L.; Seiberg, M.; Han, R. The proteinase-activated receptor-2 mediates phagocytosis in a Rho-dependent manner in human keratinocytes. J. Invest. Dermatol. 2003, 121, 529–541, doi:10.1046/j.1523-1747.2003.12427.x.

16. Correia, M.S.M.S.; Moreiras, H.; Pereira, F.J.C.F.J.C.; Neto, M.V.M. V; Festas, T.C.T.C.; Tarafder, A.K.A.K.; Ramalho, J.S.J.S.; Seabra, M.C.M.C.; Barral, D.C.D.C. Melanin Transferred to Keratinocytes Resides in Nondegradative Endocytic Compartments. J. Invest. Dermatol. 2018, 138, 637–646, doi:10.1016/j.jid.2017.09.042.

17. Tadokoro, R.; Murai, H.; Sakai, K.I.; Okui, T.; Yokota, Y.; Takahashi, Y. Melanosome transfer to keratinocyte in the chicken embryonic skin is mediated by vesicle release associated with Rho-regulated membrane blebbing. Sci. Rep. 2016, 6, 1–11, doi:10.1038/srep38277.

18. Thong, H.-Y.; Jee, S.-H.; Sun, C.-C.; Boissy, R.E. The patterns of melanosome distribution in keratinocytes of human skin as one determining factor of skin colour. Br. J. Dermatol. 2003, 149, 498–505, doi:10.1046/j.1365-2133.2003.05473.x.

19. Paul, D.; Achouri, S.; Yoon, Y.Z.; Herre, J.; Bryant, C.E.; Cicuta, P. Phagocytosis dynamics depends on target shape. Biophys. J. 2013, 105, 1143–1150, doi:10.1016/j.bpj.2013.07.036.

20. Hirota, K.; Ter, H. Endocytosis of Particle Formulations by Macrophages and Its Application to Clinical Treatment. In Molecular Regulation of Endocytosis; InTech, 2012; pp. 413–428 ISBN 978-953-51-0662-3.

21. Jaffe, A.B.; Hall, A. Rho GTPases: Biochemistry and biology. Annu. Rev. Cell Dev. Biol. 2005, 21, 247–269, doi:10.1146/annurev.cellbio.21.020604.150721.

22. Groves, E.; Dart, A.E.; Covarelli, V.; Caron, E. Molecular mechanisms of phagocytic uptake in mammalian cells. Cell. Mol. Life Sci. 2008, 65, 1957–1976, doi:10.1007/s00018-008-7578-4.

23. Kerr, M.C.; Teasdale, R.D. Defining Macropinocytosis. Traffic 2009, 10, 364–371, doi:10.1111/j.1600-0854.2009.00878.x.

24. Koivusalo, M.; Welch, C.; Hayashi, H.; Scott, C.C.; Kim, M.; Alexander, T.; Touret, N.; Hahn, K.M.; Grinstein, S. Amiloride inhibits macropinocytosis by lowering submembranous pH and preventing Rac1 and Cdc42 signaling. J. Cell Biol. 2010, 188, 547–563, doi:10.1083/jcb.200908086.

25. Wolff, K.; Konrad, K. Phagocytosis of latex beads by epidermal keratinocytes in vivo. J. Ultrasructure Res. 1972, 39, 262–280, doi:10.1016/S0022-5320(72)90022-6.

26. Potter, B.; Medenica, M. Ultramicroscopic phagocytosis of synthetic melanin by epidermal cells in vivo. J. Invest. Dermatol. 1968, 51, 300–303, doi:10.1038/jid.1968.132.

27. Vega, F.M.; Ridley, A.J. Rho GTPases in cancer cell biology. FEBS Lett. 2008, 582, 2093–2101.

28. Li, L.; Wan, T.; Wan, M.; Liu, B.; Cheng, R.; Zhang, R. The effect of the size of fluorescent dextran on its endocytic pathway. Cell Biol. Int. 2015, 39, 531–539, doi:10.1002/cbin.10424.

29. Castellano, F.; Montcourrier, P.; Chavrier, P. Membrane recruitment of Rac1 triggers phagocytosis. J. Cell Sci. 2000, 113, 2955–2961.

30. Fujii, M.; Kawai, K.; Egami, Y.; Araki, N. Dissecting the roles of Rac1 activation and deactivation in macropinocytosis using microscopic photo-manipulation. Sci. Rep. 2013, 3, 1–10, doi:10.1038/srep02385.

31. Koike, S.; Yamasaki, K.; Yamauchi, T.; Shimada-Omori, R.; Tsuchiyama, K.; Ando, H.; Aiba, S. TLR3 stimulation induces melanosome endo/phagocytosis through RHOA and CDC42 in human epidermal keratinocyte. J. Dermatol. Sci. 2019, 96, 168–177, doi:10.1016/j.jdermsci.2019.11.005.

32. Scott, G.; Leopardi, S. The cAMP signaling pathway has opposing effects on Rac and Rho in B16F10 cells: Implications for dendrite formation in melanocytic cells. Pigment Cell Res. 2003, 16, 139–148, doi:10.1034/j.1600-0749.2003.00022.x.

